# Predicting the Distribution of Serotonergic Axons: A Supercomputing Simulation of Reflected Fractional Brownian Motion in a 3D-Mouse Brain Model

**DOI:** 10.1101/2023.03.19.533385

**Authors:** Skirmantas Janušonis, Justin H. Haiman, Ralf Metzler, Thomas Vojta

**Author notes:** **Correspondence:** Skirmantas Janušonis.

## Abstract

The self-organization of the brain matrix of serotonergic axons (fibers) remains an unsolved problem in neuroscience. The regional densities of this matrix have major implications for neuroplasticity, tissue regeneration, and the understanding of mental disorders, but the trajectories of single fibers are strongly stochastic and require novel conceptual and analytical approaches. In a major extension to our previous studies, we used a supercomputing simulation to model 1000 serotonergic fibers as paths of superdiffusive fractional Brownian motion (FBM), a continuous-time stochastic process. The fibers produced long walks in a complex, threedimensional shape based on the mouse brain and reflected at the outer (pial) and inner (ventricular) boundaries. The resultant regional densities were compared to the actual fiber densities in the corresponding neuroanatomically-defined regions. The relative densities showed strong qualitative similarities in the forebrain and midbrain, demonstrating the predictive potential of stochastic modeling in this system. The current simulation does not respect tissue heterogeneities, but can be further improved with novel models of multifractional FBM. The study demonstrates that serotonergic fiber densities can be strongly influenced by the geometry of the brain, with implications for brain development, plasticity, and evolution.

## 1. INTRODUCTION

The self-organization of the brain serotonergic matrix remains an unsolved problem in neuroscience. This matrix, or a meshwork of axons (fibers), is present in virtually all brain regions, likely across the entire vertebrate clade (Wolters et al., 1985; Stuesse et al., 1991; Metzger et al., 2002; López and González, 2014; Janušonis, 2018; Awasthi et al., 2020). It appears to be fundamentally associated with neuroplasticity (Lesch and Waider, 2012; Daws et al., 2022), raising new questions (Vargas et al., 2023), and is remarkable in its ability to robustly regenerate in the adult mammalian brain (Jin et al., 2016; Cooke et al., 2022). In mammals, serotonergic axons originate exclusively in the brainstem raphe nuclei; the total number of serotonin (5-hydroxytryptamine, 5-HT)-synthesizing neurons has been estimated at 20,000-26,000 in mice and rats and at around 450,000 in humans (Jacobs and Azmitia, 1992; Okaty et al., 2019). Comprehensive analyses have shown that this population of neurons is extremely diverse in its transcriptomes (Okaty et al., 2019; Ren et al., 2019; Okaty et al., 2020).

Recent studies have led to an intriguing picture of this system. On the one hand, state-of-the-art methods, such as single-cell RNA-seq and viral-genetic tract-tracing, have revealed its deep, *deterministic-like* organization. In particular, specific transcriptome profiles (including neurotransmitter complements and excitability signatures) have been mapped to nested neuron clusters, each with its anatomically-defined set of projections (Ren et al., 2018; Okaty et al., 2019; Ren et al., 2019; Okaty et al., 2020). On the other hand, serotonergic axons have been shown to be strongly *stochastic* in their trajectories. These trajectories can be viewed as unique paths (realizations) of rigorously-defined spatial stochastic processes (Janušonis and Detering, 2019; Janušonis et al., 2019; Janušonis et al., 2020; Vojta et al., 2020). Both approaches are innovative in that they seek to reveal how precisely-specified events at the single-neuron level result in the self-organization of the entire serotonergic system, in space and time. Their conceptual underpinnings can be traced back several decades (Katz et al., 1984; Waterhouse et al., 1986).

The emerging duality of the serotonergic system, with well-defined deterministic and stochastic components, may reflect the fundamental principles of the self-assembly of neural tissue. The constructive interplay between dynamic determinism and stochasticity has been suggested by studies that have used various experimental platforms. An individual serotonergic neuron may express a specific gene network, distinctly different from other “adjacent” networks, but it may still be able to perform switch-like transitions among them, in the presence of environmental “noise” (Okaty et al., 2019; Ren et al., 2019). Dorsal raphe serotonergic neurons vary in their firing rates, but this variability may be captured by a single normal distribution, across different anatomical locations (Mlinar et al., 2016). Several classes of serotonergic axons have been defined based on their morphology (Kosofsky and Molliver, 1987), but recent studies have demonstrated that axonal morphology may undergo significant changes in a regionally- and developmentally-dependent manner (Gagnon and Parent, 2014; Maddaloni et al., 2017; Nazzi et al., 2019; Okaty et al., 2019; Andersson et al., 2020). In primary cell cultures, adjacent axon segments of serotonergic neurons can strongly vary in their caliber, varicosity size, and other features (Hingorani et al., 2022). This variability is likely partially stochastic, as a consequence of the strongly stochastic properties of neural tissue at the microscopic level (Jang et al., 2010; Nicholson and Hrabetova, 2017; Hrabetova et al., 2018; Hingorani et al., 2022). Furthermore, different neuron clusters may show preference for different targets; however, individual serotonergic axons within the same cluster may differ in their collateralization and target specificity (Gagnon and Parent, 2014; Ren et al., 2019). Also, serotonergic neurons that send their axons to the same anatomical region may not be physically clustered (Okaty et al., 2019). In experimental brain injury models, regenerating serotonergic fibers do not follow their previous trajectories and create new tortuous paths (Jin et al., 2016).

This study sought to investigate the potential of the purely stochastic component of serotonergic axons (fibers), with regard to the self-organization of regional serotonergic densities. Individual serotonergic fibers were modeled as paths of fractional Brownian motion (FBM), a continuous-time stochastic process (Mandelbrot and Van Ness, 1968; Biagini et al., 2010). FBM generalizes normal Brownian motion (BM) in that it allows positive and negative correlations among displacement increments (in BM, used to describe free diffusion of particles, non-overlapping displacements are assumed to be statistically independent). FBM process is parametrized with the Hurst index (0 < *H* < 1), which leads to three distinctly different regimes: subdiffusion (*H* < 0.5), BM (*H* = 0.5), and superdiffusion (*H* > 0.5). We have previously shown that serotonergic fiber trajectories can be modeled with superdiffusive FBM (*H* ≈ 0.8) (Janušonis et al., 2020).

In a previous study, we have analyzed the distributions of FBM-driven fibers in a selected set of two-dimensional (2D) shapes based on coronal sections of the adult mouse brain (Janušonis et al., 2020). However, the long-range dependence among spatial displacements implies that the trajectory of a fiber at a given coronal level depends on its history in the threedimensional (3D) space (*e.g*., at more rostral or caudal coronal sections). In this study, we performed a supercomputing simulation of a large number of fibers in a complex 3D-shape that was constructed from a serially sectioned mouse brain. The simulation was based on reflected (boundary-limited) FBM (rFBM), the theoretical properties of which have been investigated by our group (Wada and Vojta, 2018; Guggenberger et al., 2019; Vojta et al., 2020). It is a major extension of the previous study in that the simulation captured the entire 3D-geometry of the brain, virtually eliminating the dependence of the results on the sectioning plane.

In the adult mouse brain, this geometry contains some elaborate elements (*e.g.*, the folded hippocampus) and is enriched with many dense, fully developed axon tracts. These tracts (*e.g.*, the anterior commissure, the corpus callosum, the fasciculus retroflexus) are nearly impermeable to serotonergic axons and act as obstacles in simulations (Janušonis et al., 2020). To reduce this complexity, a late-embryonic mouse brain (at embryonic day (E) 17.5) was used to construct the 3D-shape. This selection is further justified by the evidence that serotonergic neurons mature early; by this age, their axons are already present in the telencephalon and reach the cortical plate in mice and rats (Wallace and Lauder, 1983; Brüning et al., 1997; Janušonis et al., 2004). The geometries (*e.g.*, curvatures, distances) of the embryonic and adult brain are not identical, but they share the same fundamental topology and major features. Strictly speaking, neither the embryonic brain nor the adult brain is the “correct” static shape in this context: a developmentally accurate approach would require simulating fiber trajectories as the shape itself increases in size and morphs. However, without accurate experimental information about the relative dynamics of both processes such simulations are unlikely to produce robust results. State-of-the-art imaging techniques have provided new insights into the growth dynamics of single serotonergic axons (Jin et al., 2016; Hingorani et al., 2022), but live-imaging of serotonergic fibers in the intact developing brain requires further technological advances.

The verification of simulation results requires accurate information about actual serotonergic fiber densities. These densities have been a major focus of investigation since the discovery of 5-HT-producing neurons in the brain (Hokfelt, 2016). Their regionally-specific estimates, independent of the neuroanatomical origin of the fibers, initially relied on the detection of 5-HT or the serotonin transporter (SERT) (Steinbusch, 1981; Lidov and Molliver, 1982; Foote and Morrison, 1984; Voigt and de Lima, 1991; Way et al., 2007; Vertes et al., 2010; Linley et al., 2013; Belmer et al., 2017). These markers are specific to serotonergic neurons and strongly overlap (Belmer et al., 2019), with some caveats (Lebrand et al., 1996; Lebrand et al., 2006; Maddaloni et al., 2017). However, they are directly associated with local 5-HT accumulation and release, and therefore may not visualize fibers or their segments with low 5-HT levels (*e.g*., if they have a fine caliber and contain no varicosities). The 5-HT signal can show striking variability along the path of a single fiber in cell-culture preparations (Hingorani et al., 2022). In addition, the relative abundance of SERT may depend on the local diffusivity of extracellular 5-HT, which in turn depends on the local properties of the extracellular space (Syková and Nicholson, 2008; Hrabetova et al., 2018; Okaty et al., 2019). Furthermore, serotonergic neurons can release other neurotransmitters, such as glutamate, and their terminals can segregate by the preferred neurotransmitter (Okaty et al., 2019).

In the several past decades, studies have mapped the projections of specific raphe nuclei, often using sensitive visualization procedures (*e.g*., efficient neural tract tracers combined with immunoperoxidase detection) (Vertes, 1991; Morin and Meyer-Bernstein, 1999; Vertes et al., 1999). These connectomics-driven approaches are independent of 5-HT accumulation, but the obtained densities reflect only the contribution of the selected nucleus. In addition, they strongly depend on the uptake efficiency of the tracer (*i.e*., some serotonergic neurons and their axons are likely to remain unlabeled).

More recently, transgenic mouse models have allowed extremely accurate visualization of all serotonergic axons, irrespective of their anatomical origin or 5-HT content. For example, a fluorescent reporter (*e.g*., EGFP) can be expressed under the promoter a serotonergic neuron-specific gene (*e.g*., *Tph2*) and further enhanced with immunohistochemistry. Since reporter proteins can be transported to distal axon segments, they can reveal the dynamics and susceptibility of serotonergic fibers with unprecedented precision (Maddaloni et al., 2017; Maddaloni et al., 2018). However, these studies have focused on targeted brain regions, due to the complexity of single-fiber analyses.

These research trends have resulted in a paradoxical lack of modern, comprehensive atlases of serotonergic fiber densities. Such atlases should (1) cover the *entire* brain and (2) detect *all* physically present serotonergic fibers – with high specificity and sensitivity, irrespective of their signaling state (*e.g*., 5-HT content). The recent publication of a detailed topographical map of the serotonergic fiber densities in the entire adult mouse brain has now partially filled this knowledge gap (Awasthi et al., 2020). This map, based on the expression of EGFP under the SERT gene promoter, currently does not include the adult female mouse brain or developmental stages.

In summary, this study brings together several recent developments: the models of serotonergic fibers as rFBM-paths, the advances in the theory of rFBM, and the new comprehensive map of serotonergic fiber densities. By using a supercomputing simulation, we attempted to predict regional fiber densities based purely on the geometry of the brain and compared them to the neuroanatomical data.

## 2. METHODS

### 2.1. Brain Sections

A timed-pregnant mouse (Charles River Laboratories) was deeply anesthetized at E17.5 with a mixture of ketamine (200 mg/kg) and xylazine (20 mg/kg), and the embryos were dissected into cold 0.1 M phosphate-buffered saline (PBS, pH 7.2). They were immediately decapitated, and their brains were dissected with fine forceps under a stereoscope. Two embryonic brains were rinsed in PBS and immersion-fixed in 4% paraformaldehyde overnight at 4°C. They were cryoprotected in 30% sucrose for two days at 4°C and embedded in 20% gelatin (type A) in a Peel-A-Way mold, with an insect pin pushed through the mold in the rostro-caudal orientation just dorsal to the brain. After one hour at 4°C, the gelatin block was removed, trimmed, and incubated for 3 hours in formalin with 20% sucrose at room temperature. It was sectioned coronally from the olfactory bulbs to the caudal midbrain at 40 μm thickness on a freezing microtome into 96-well trays with PBS. The lower brainstem and the cerebellum were not included. In order to avoid distance distortions in the rostro-caudal axis, empty wells were used to mark damaged or missing sections. Every other section was mounted onto gelatin/chromium-subbed glass slides, allowed to air-dry, and imaged uncoverslipped on a Zeiss Axio Imager Z1 in bright field with a 1× objective (NA = 0.025) (Fig. 1A). All animal procedures have been approved by the UCSB Institutional Animal Care and Use Committee.

**Figure 1.**
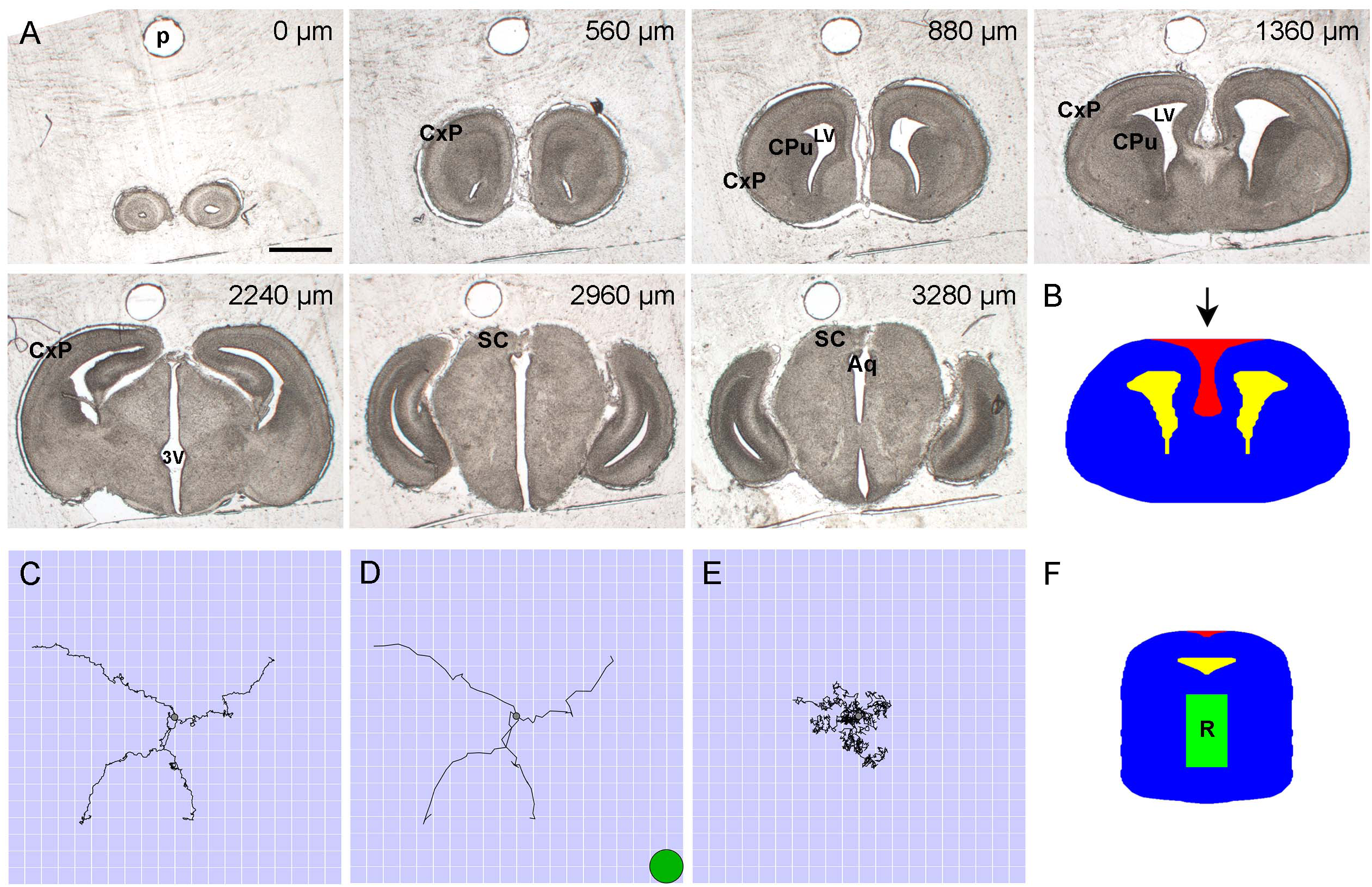
(**A**) Representative bright-field images of unstained and uncoverslipped coronal brain sections (E17.5) used in the set. The rostrocaudal distances are shown at the top of the panels. In the simulation, tissue non-homogeneities were disregarded. Aq, cerebral aqueduct; CxP, cortical plate; CPu, caudate-putamen; LV, lateral ventricle; SC, superior colliculus; 3V, third ventricle. Scale bar = 1 mm. (**B**) Each of the constitutive shapes in a section was fully described by its leftmost and rightmost points at each dorsoventral level. The 2D-shape represents the section at the rostrocaudal distance of 1360 μm. The allowed region is shown in blue; the forbidden regions are shown in yellow and red (the lateral ventricles and the dorsomedially oriented concavity, respectively). (**C**) Four sample FBM (*H* = 0.8) trajectories with mean zero and *σ* = 0.4 spatial units, from 0 to 20 time-units with the walk-step of 0.05 time-units. The side of one cell is one spatial unit (corresponding to 6.6 μm in the physical brain). All trajectories start at the center (gray circle) of the region containing 20 × 20 grid cells. The small step is used to show the fine details of the process. (**D**) The same trajectories shown with the walk-step of one time-unit (as actually used in the simulation). The green circle shows the relative size of a typical neuronal soma in the adult brain (with the diameter of around 13 μm). (**E**) Four sample trajectories of normal Brownian motion (*H* = 0.5) with mean zero and *σ* = 0.4 spatial units, from 0 to 20 time-units with the walk-step of 0.05 time-units. The trajectories are shown for comparison (normal Brownian motion was not used in the simulation; it produces trajectories with uncorrelated increments). (**F**) All fibers started in the rostral raphe region (R), approximated by a cuboid under the aqueduct.

### 2.2. Conversion of Images to a Stack of 2D-Shapes for Simulations

The section series of both brains were examined, and one series was selected for further processing. The sections were aligned in the rostro-caudal axis in Reconstruct (SynapseWeb), as shown in our previous publication (Flood et al., 2012). The pin-hole in the gelatin was used as a fiducial marker (Fig. 1A). The aligned images were imported in Photoshop 23 (Adobe, Inc.), and brain contours were outlined by an expert neuroanatomist using the magnetic or polygonal lasso tools. The contours were converted into binary images (white shapes on a black background). If a section contained more than one shape (*e.g*., the outer outline and ventricular spaces), they were saved as separate images, in the same aligned geometric space. In order to reduce the complexity of the geometry, the folding of the hippocampus was not respected (*i.e*., the hippocampus was represented by a medial pallial region with no internal structure).

The binary images were imported into Wolfram Mathematica 13 (Wolfram Research, Inc.) and processed as previously described (Janušonis et al., 2020). Briefly, each closed contour was converted into an ordered set of points, smoothed, bilaterally symmetrized, and transformed into a different format represented by an *N* × 3 matrix (where *N* is the number of the rows). The rows represented the consecutive *y*-coordinates (with no gaps), from the most dorsal level to the most ventral level of the contour. Each row contained three values: a *y*-coordinate and the leftmost and rightmost *x*-coordinates of the contour at this *y*-coordinate (both *x*- and *y*-coordinates were integers). Since this format cannot capture concavities oriented in the dorsoventral direction, they were coded as separate contours representing “forbidden” regions (in addition to the ventricular spaces) (Fig. 1B). In this two-dimensional (2D)-integer grid, the side of each square cell represented the physical distance of 6.6 μm in the physical brain.

Next, a three-dimensional brain model was built from the section stack. First, missing or damaged sections were recreated by linear interpolation between neighboring sections to maintain even (80 μm) rostro-caudal steps between any two consecutive sections. This process was guided by the known anatomy, and the resulting interpolated sections were checked manually for anatomical correctness. Second, in order to produce a three-dimensional simulation grid, each section was further subdivided into a set of twelve virtual sections of around 6.6 μm in thickness using linear interpolation. The resulting simulation grid cells (voxels) were thus cubes whose linear size corresponded to 6.6 μm in the physical brain.

### 2.3. Supercomputing Simulations

The fiber densities were produced by 1000 simulated fibers. Each fiber was represented by the path of a discrete three-dimensional FBM (Qian, 2003). Specifically, the trajectory of the random walker moved according to the recursion relation *r*_*n*+1_ = *r_n_* + *ξ_n_*, where *r_n_* is the (threedimensional) walker position, and the steps (increments) *ξ_n_* are a three-component discrete fractional Gaussian noise (Fig. 1C, D). The statistically-independent *x, y*, and *z* components of *ξ_n_* were Gaussian random numbers with mean zero and variance *σ*^2^, and each component had long-range correlations between steps (in contrast to normal Brownian motion (Fig. 1E)). The corresponding noise covariance function between steps *m* and *m* + *n* was given by 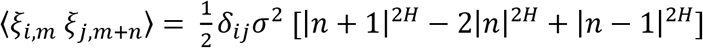, where *H* is the Hurst index, *δ_ij_* is the Kronecker delta, and *i, j* = *x*, *y*, *z* denotes the three space dimensions. The Fourier-filtering method (Makse et al., 1996) was employed to generate these long-range correlated, stationary random numbers on the computer (Wada and Vojta, 2018; Vojta et al., 2020). This efficient method allowed us to create particularly long trajectories. The main simulations were performed for Hurst index *H* = 0.8, but we also tested other *H* values. The root mean-square step size was set to *σ* = 0.4 grid units, corresponding to 2.6 μm in the physical brain (smaller than the diameter of a single neuron). All paths started in the rostral raphe region, given by a cuboid with the approximate physical dimensions 680 × 1200 × 400 μm^3^ in the mediolateral, dorsoventral, and rostrocaudal directions, respectively (Fig. 1F). Each trajectory consisted of 2^25^ ≈ 33.6 million walk-steps (each 1 time-unit). The length of the trajectories was sufficient for the relative densities to reach a steady state.

If the propagating fiber encountered a boundary (*i.e*., an outer or inner contour), it was reflected. Our previous extensive analyses have shown that the choice of the reflection condition has virtually no effect on the simulation results, with the exception of a very narrow region (a few steps wide) at the boundary (Vojta et al., 2020). Therefore, the following simulation used the simple condition under which a step that would move the leading fiber end into the forbidden region was simply not carried out. Deciding whether a given point is inside or outside of a complex three-dimensional shape is a complicated problem in computational geometry. Our approach to modeling the geometry in terms of virtual sections and boundary contours within each section (as described above) allowed us to implement an efficient local inside-outside test. The most rostral and caudal sections of the model brain geometry were treated as reflecting boundaries in the rostro-caudal direction.

After the simulation, the resultant densities were evaluated in non-overlapping cubes (composed of 2 × 2 × 2 grid cells, to suppress noise and achieve more robust estimates). The local density (*d_s_*) was determined by counting the total number of random-walk segments inside each cell. These local densities were normalized to the total sum of one in the entire 3D-brain volume, to remove any dependence of the results on the arbitrarily chosen trajectory length. To facilitate comparisons between the simulated fiber densities and published densities (typically, in immunostained sections), the raw simulation densities were transformed to “optical densities” using a Beer-Lambert law-like transformation *d_o_* = 1 – exp(–*kd_s_*) (Janušonis et al., 2020), with an empirically optimized *k* value (*k* = 10^8.3^). This transformation constrained density values to a finite interval (from zero to one). They were used for all further analyses and figures. The graphical density maps were produced in Mathematica 13, using the “rainbow” and “heat” color schemes.

All supercomputing simulations were written in Fortran and carried out on the Pegasus cluster at the Missouri University of Science & Technology, using parallel processing on several hundred CPU cores.

### 2.4. The evaluation of simulated fiber densities

The simulated fiber densities were evaluated using a comprehensive map of the serotonergic fiber densities in the adult male mouse brain (on the C57BL/6J background) (Awasthi et al., 2020) and an atlas of the developing mouse brain (Paxinos et al., 2020).

## 3. RESULTS

The fiber densities obtained in the simulation are shown in Figure 2. For accurate quantitative comparisons, the density values along selected cuts (1D-segments) are included (Fig. 3).

**Figure 2.**
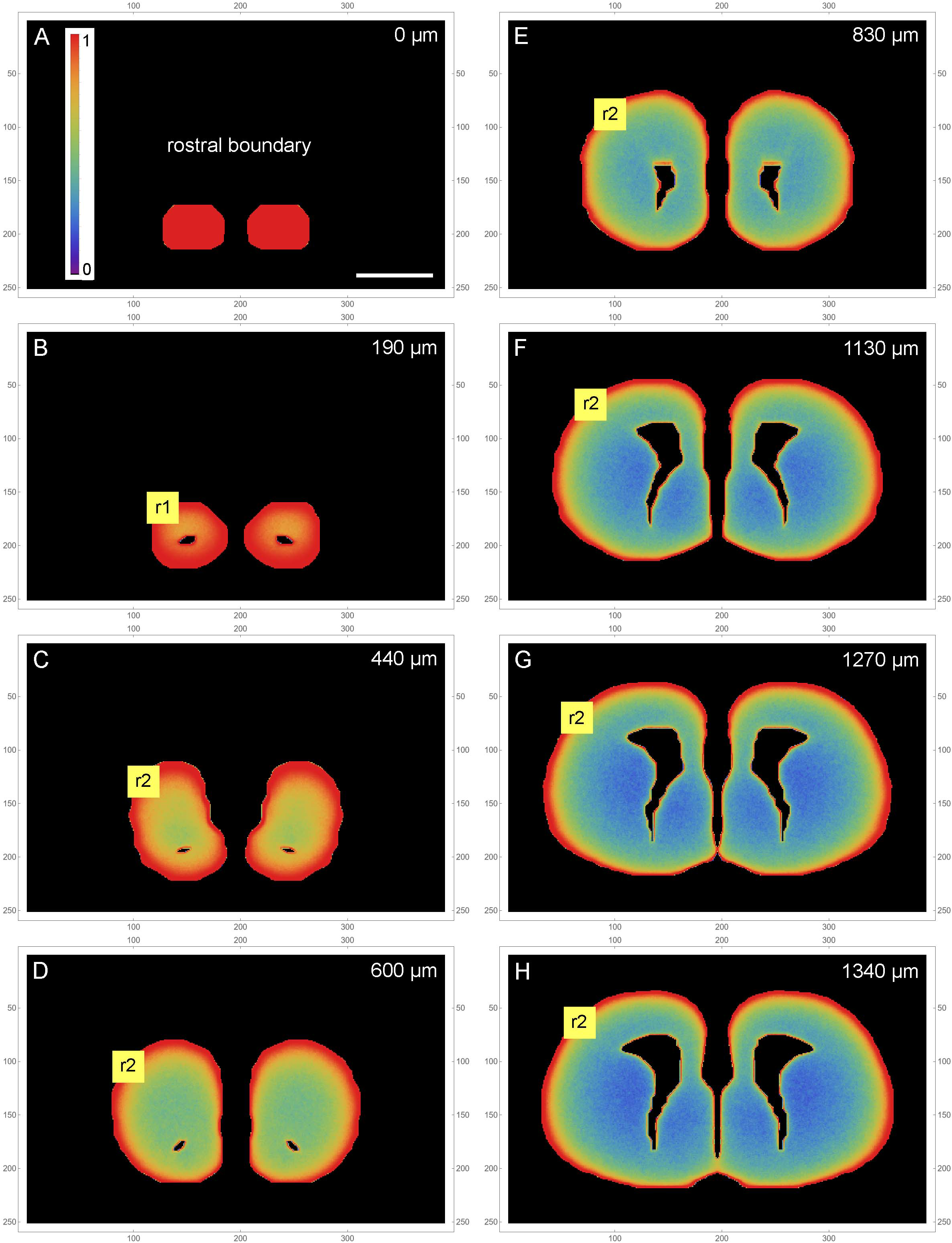
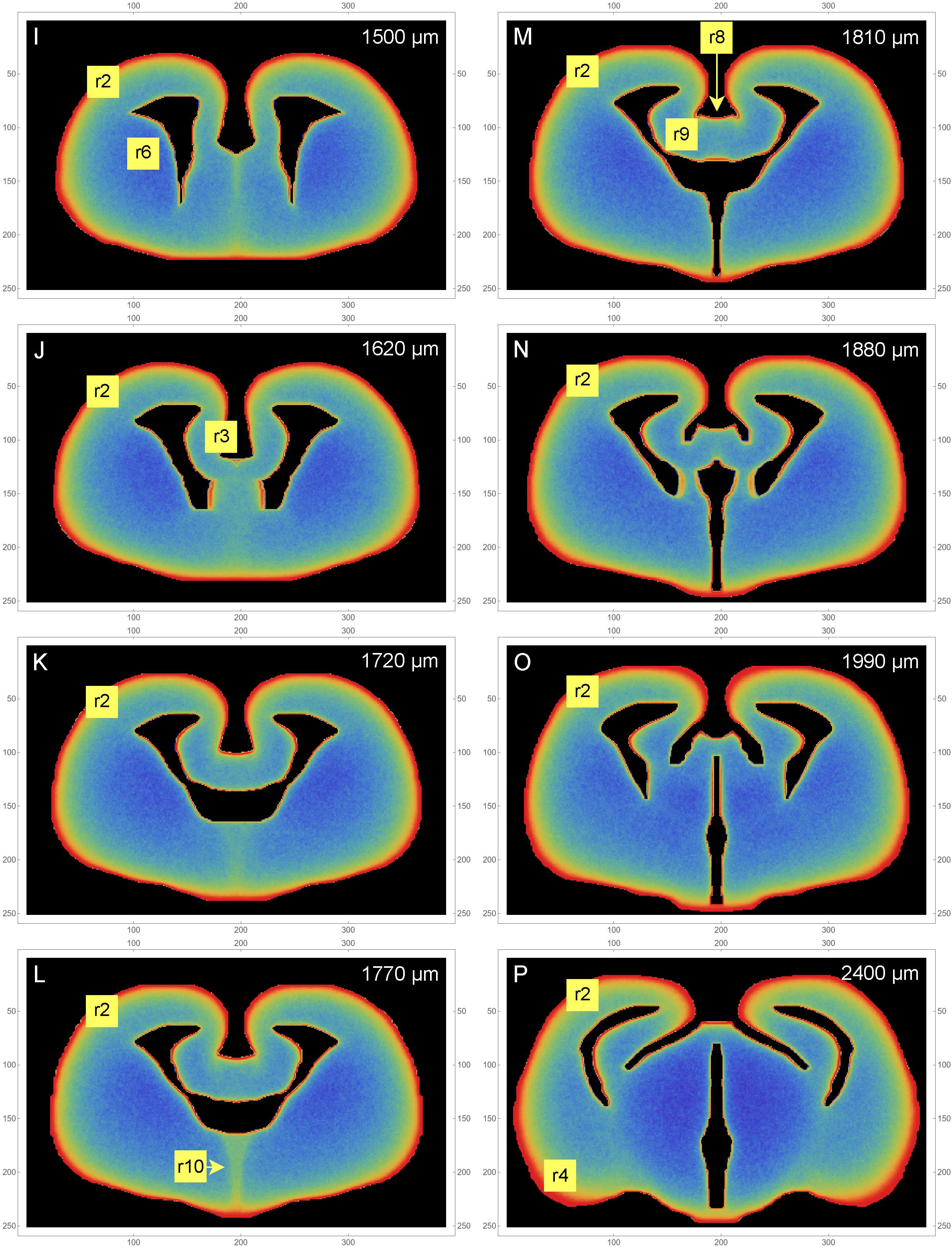
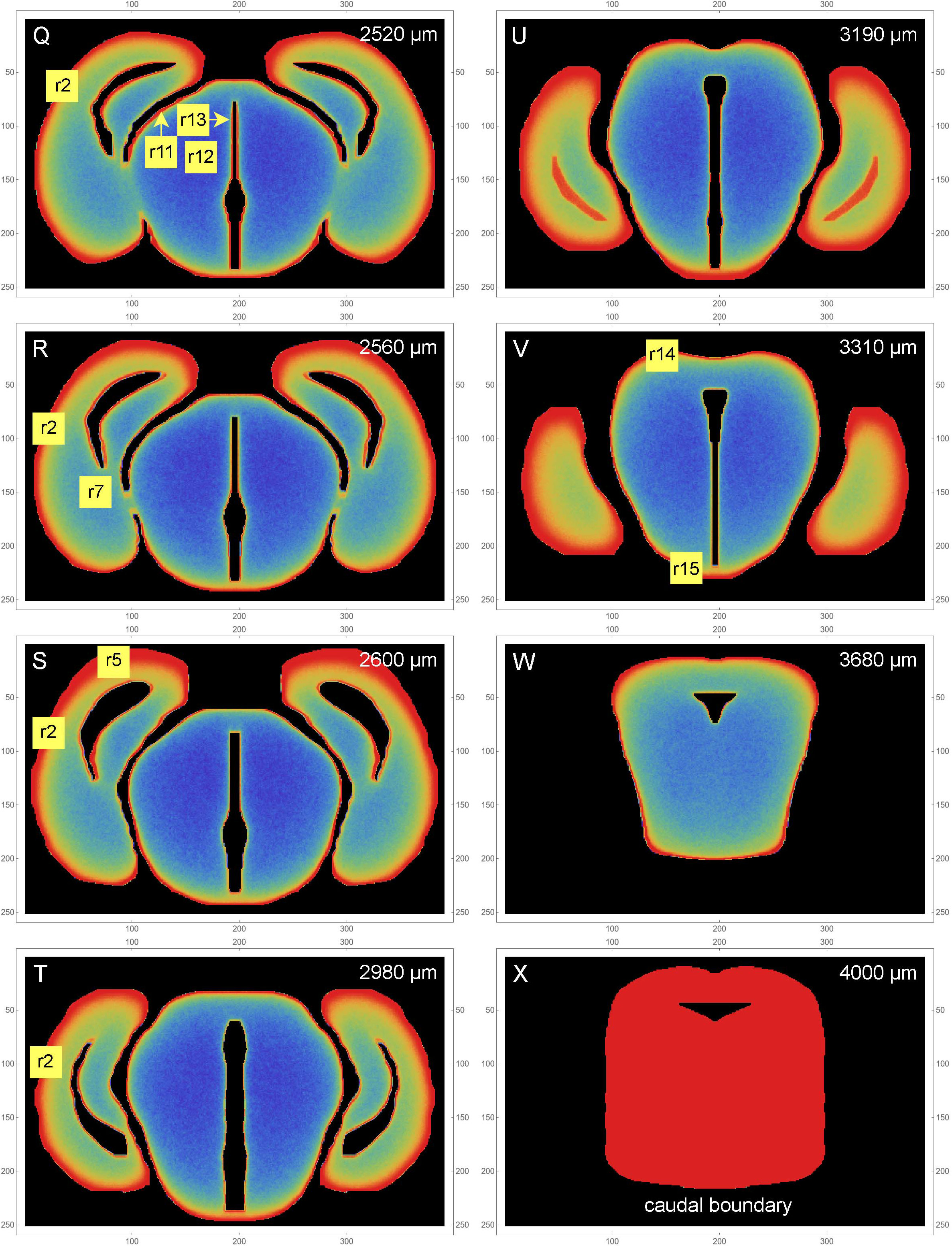
(*in 3 parts*) A color-map atlas of the simulated fiber densities, from the rostral brain pole to the most caudal section in the set. Low densities are blue, medium densities are green, and high densities are red. Key rostrocaudal levels and transitions are shown (note that the rostrocaudal distances between panels vary; they are shown at the top of the panels). The selected regions (r1-r15) are discussed in the Results section. The frame numbers indicate square cells after the 2 × 2 pooling. Scale bar = 1 mm.

**Figure 3.**
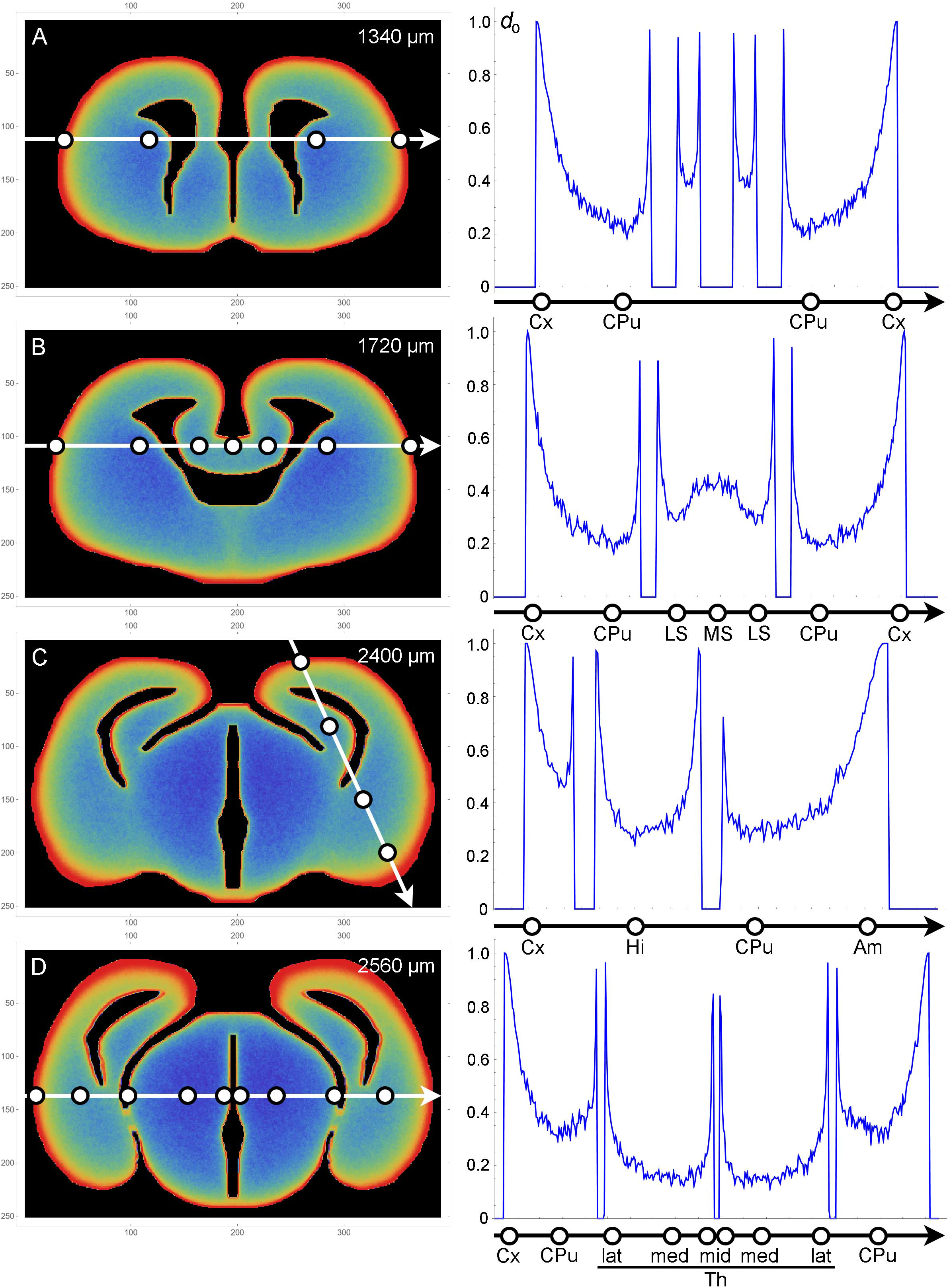
Plots of fiber densities (right) along selected one-dimensional cuts across four sections (left). Regions important for comparisons with the actual serotonergic fiber densities are marked with circles. Am, amygdala region; CPu, caudate-putamen; Cx, cortex; Hi, hippocampal (medial pallial) region; LS, lateral septum; MS, medial septum; Th, thalamus (lat, lateral; med, medial; mid, midline); *d*_o_, optical density (0-1). The frame numbers indicate square cells after the 2 × 2 pooling.

The extremely high accumulation of fibers in the most rostral section (Fig. 2A) reflects its position as a coronal boundary of the 3D-shape (*i.e*., fibers cannot advance rostrally beyond it). The actual olfactory bulb extends a little further, but this section can also be considered to be the most superficial *layer* of the rostral pole.

Generally, the simulated fibers produced the highest fiber densities close to the tissue borders (the pia as the outer border and the ependyma of the ventricular system as the inner border). However, the memory (autocorrelation) of the fibers and the complex shape they interacted with led to other density variations that could not be predicted purely from the local geometry. We next discuss the key findings and compare them to the experimentally established densities of serotonergic fibers in the adult mouse brain, based on the comprehensive map of Awasthi et al. (2020). In this map, regional densities (*d*) are semi-quantitatively evaluated on the scale of 0 (extremely low) to 6 (extremely high).

In the olfactory bulb, the dense accumulation of the simulated fibers at the outer border (Fig. 2B [r1]) corresponds to the glomerular layer, the layer with the highest serotonergic fiber density in this structure (*d* = 5). In the telencephalon proper, the neural tissue at the outer and inner borders corresponds to the adult cortical layer I and periventricular regions, respectively. In most regions, the high-density band at the outer border (r2) was considerably thicker than the bands at the inner borders (Fig. 2C-T). This pattern was in register with the high densities (*d* = 5) of serotonergic fibers in layer I of virtually all, functionally different adult cortical regions (in which no other layer exceeds the layer I density). These regions cover the entire rostro-caudal and medio-lateral extents and include the prefrontal cortex, the motor cortex, the somatosensory cortex, the auditory cortex, and the piriform cortex, with the minor exception of the retrosplenial cortex (RSC; *d* = 3). However, the RSC (areas A29c and A30 of the cingulate cortex (Vogt and Paxinos, 2014)) has an extensive, prominently flattened cortical region (A29c), where the two hemispheres press against each other at the midline. The exact boundaries of A29c are not readily identifiable in the embryonic mouse brain (Paxinos et al., 2020), but in the simulation similar flat cingulate regions showed narrower high-density bands (Fig. 2J [r3]). The basolateral amygdala (BLA) has an exceptionally high density of serotonergic fibers (*d* = 5-6), with the adjacent regions showing a similar pattern (the basomedial amygdala (BMA) and the piriform cortex: *d* = 3-5; the endopiriform nucleus: *d* =5-6). The simulated fibers produced a particularly thick high-density band in the corresponding region (Fig. 2P [r4], Fig. 3C). However, the simulated fibers did not show the tendency to decrease in density in cortical regions in the rostro-caudal direction, as reported by Awasthi et al. (2020), and produced another thick high-density band in a region corresponding to the future visual cortex (Fig. 2S [r5]). This discrepancy may reflect limitations of the model, but it can also be due to other causes: in the adult brain the visual cortex extends more caudally (to the cerebellum); serotonergic neurons that reach the most caudal cortical regions incur an energetic cost in supporting extremely long axons (with no such limitation in simulations); and the detection of distal axonal segments can be affected by the long-distance transport of EGFP, the protein used for fiber visualization.

The simulated fibers produced a low density in the region corresponding to the caudate-putamen (Fig. 2I [r6], Fig. 3A-B), despite its proximity to the medially-bulging edge of the lateral ventricles. The fiber density increased in the more caudal caudate-putamen regions (Fig. 2R [r7], Fig. 3A-C), generally consistent with the low-to-moderate (*d* = 2-4) densities of serotonergic fibers in this region, with the same gradient.

The complex geometry of the septal region led to a subtly higher density of simulated fibers in the medial septum than the lateral septum (Fig. 2M [r8, r9], Fig. 3B). With the exception of the rostral part of the lateral septum, the brain shows a similar gradient of serotonergic fiber densities (the medial septum: *d* = 3-5; the lateral septum: *d* = 1-3).

Just rostral to the transition to the third ventricle, simulated fibers produced a distinct median band with an elevated fiber density (Fig. 2L [r10]). This region corresponds to the region of the preoptic hypothalamus, which has a high density of serotonergic fibers (e.g., *d* = 5 in the median preoptic nucleus and parts of the medial preoptic area). The higher density was induced by the location of this region in the 3D-space. Specifically, it represents a coronal boundary to medially located fibers that move caudally: they cannot enter the third ventricle at the adjacent caudal levels and are “reflected,” with an accumulation. This same geometry may be a contributing factor in the actual brain.

In the diencephalon and mesencephalon, the highest serotonergic fiber densities tend to be located near the outer tissue border or around the ventricular spaces (see Figure 5 of Awasthi et al. (2020)). This general pattern is strongly consistent with the density distributions produced by the simulated fibers (Fig. 2O-W). In the actual brain, it can be interrupted by major axon fascicles (*e.g*., the cerebral peduncle) that cannot be easily penetrated by individual serotonergic fibers (Awasthi et al., 2020; Janušonis et al., 2020). The thalamic lateral posterior (LP), lateral dorsal (LD), and lateral geniculate (LG) nuclei, all contributing to the dorsal outer border of the thalamus, have high serotonergic fiber densities (*d* = 4-5, *d* = 4-5, *d* = 6, respectively). The simulated fibers also produced high densities in this region (Fig. 2Q [r11], Fig. 3D). Notably, in the adult mouse brain the LG has the highest outer curvature. In contrast, the deeper thalamic nuclei, such as the ventral posteromedial and posterolateral nuclei (VPM and VPL), the ventromedial nucleus (VM), the posterior nucleus (PO), and the deep part of the medial geniculate nucleus (MG), have serotonergic fiber densities that are among the lowest in the entire brain (*d* = 2 in the MG and *d* < 1 in the rest of the set). The simulated fibers also produced especially low densities in this region (Fig. 2Q [r12], Fig. 3D). The thalamic midline nuclei, located close to the third ventricle, again have high serotonergic fiber densities (*d* = 6), consistent with the accumulation of the simulated fibers at inner borders (Fig. 2Q [r13], Fig. 3D).

The mesencephalon followed a similar general pattern. The simulated fibers produced the highest densities in the regions of the superior colliculus (SC) (Fig. 2V [r14]) and the broadly defined ventral tegmental region (Fig. 2V [r15]), both of which have high serotonergic fiber densities (Awasthi et al., 2020). Despite this consistency, caution should be exercised at this coronal level because it is close to the origin of the serotonergic fibers (emerging from the raphe nuclei under the cerebral aqueduct) and it also contains several major axon fascicles deep in the tissue. Also, the caudal mesencephalon represents an unnatural boundary of the simulated central nervous system (Fig. 2X), which in reality is given by the caudal end of the spinal cord.

A selected subset of the sections is shown in pseudo-monochrome, to support direct comparisons with densities visualized with immunohistochemical procedures (Fig. 4).

**Figure 4.**
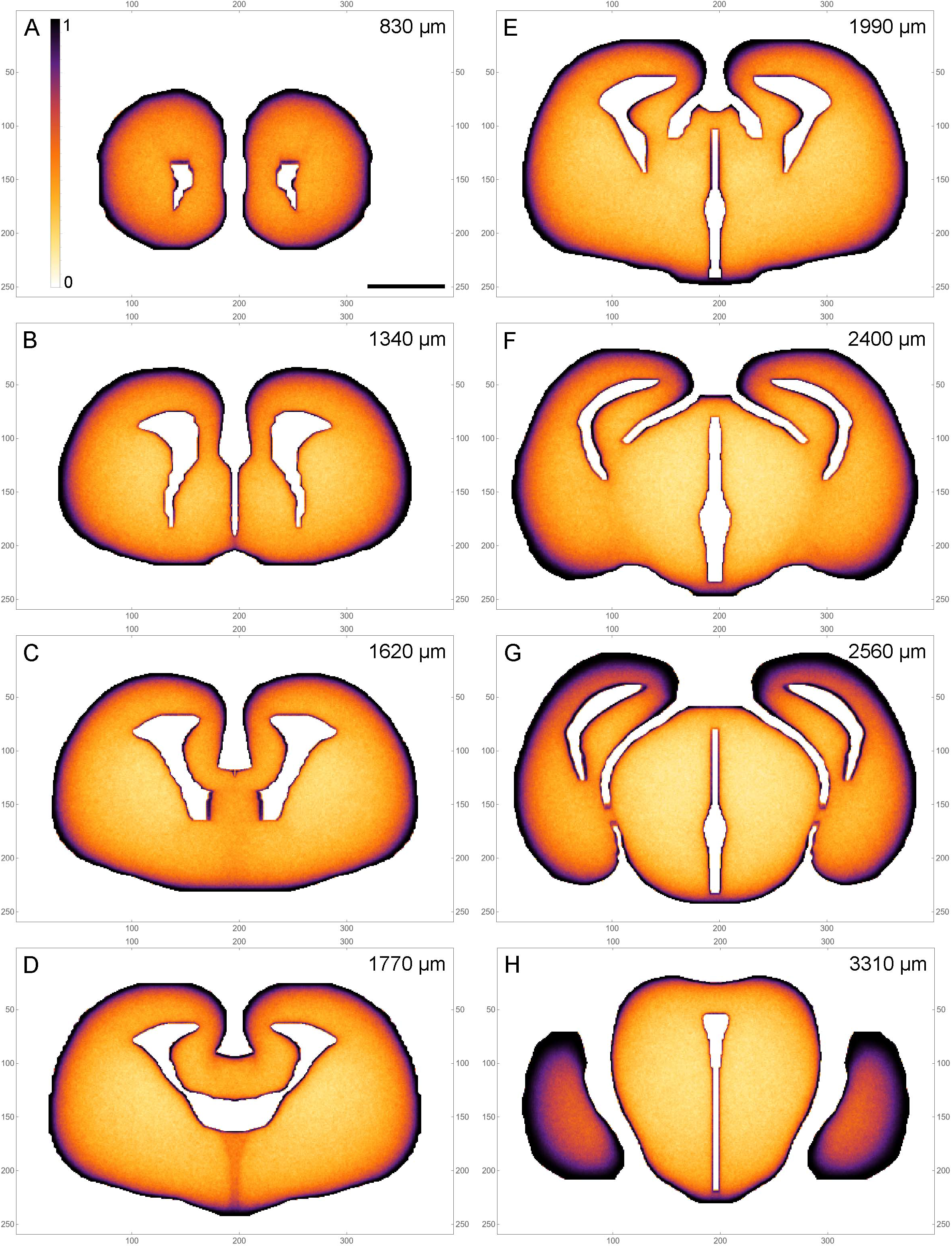
A pseudo-monochrome map atlas of the simulated fiber densities (a subset of the set in Figure 2). The colors simulate immunohistochemical visualization with 3,3’-diamonobenzidine (DAB); low densities are light, high densities are dark. The rostrocaudal distances are shown at the top of the panels. The frame numbers indicate square cells after the 2 × 2 pooling. Scale bar = 1 mm.

## 4. DISCUSSION

The supercomputing simulation produced a predictive map of serotonergic fiber densities, based on the model of fibers as rFBM paths. In addition to its contribution to neuroscience, this work is the first simulation of rFBM in any complex 3D-shape, with potential applications in other fields (Vojta et al., 2020).

The obtained densities depended only on the fundamental properties of rFBM as a stochastic process and the geometry of the 3D-shape (including the ventricular spaces). Despite this conceptual simplicity, the simulated densities approximated the relative intensities of serotonergic fiber densities in many brain regions. Exact quantitative comparisons are currently infeasible because regional serotonergic densities are traditionally reported only descriptively or as observer scores limited to a small set of discrete values (*e.g*., 0-6 in Awasthi et al. (2020)). This approach is partially motivated by the need to combine the overall signal intensity with the morphology of individual fibers; for example, the same overall signal intensity can be produced by a high density of fine-caliber fibers or by a lower density of larger-caliber axons. As a consequence, the semi-quantitative scores are assigned by expert viewers but can still be influenced by subjective biases. In contrast, assessments of relative densities, especially in adjacent or similar brain regions, are likely to be highly accurate. Therefore, we followed the same approach in our comparisons, which were nearly independent of the shape of the monotonic function used to transform raw simulated fiber densities to the corresponding (immunohistochemistry-like) signal intensities.

As expected from our previous simulations in 2D-shapes, high fiber densities were produced near the borders of the 3D-shape (Janušonis et al., 2020). Elevated serotonergic fiber densities have been reported in many of the corresponding brain regions, despite major differences in their functional roles (Awasthi et al., 2020). These regions include the most superficial layer (layer I) of virtually all cortical areas, the thalamic nuclei at the lateral and medial (ventricular) borders, the superior colliculus, the hypothalamus, the ventral region of the tegmentum, and other major neuroanatomical structures (Figs. 2–4). The local border curvature and the geometry of adjacent regions further contributed to the intensity and gradients of fiber densities. In particular, this effect produced a high-density region in the area corresponding to the amygdala complex. This region extended deep in the tissue (Fig. 3C), consistent with the actual serotonergic fiber densities in the brain (Awasthi et al., 2020). Generally, high serotonergic fiber densities in these regions also have been reported in other mapping studies (Vertes, 1991; Morin and Meyer-Bernstein, 1999; Vertes et al., 1999; Vertes et al., 2010; Linley et al., 2013), with some variability due to the technical limitations discussed in the Introduction (*e.g*., the targeting of specific raphe nuclei). Compared to the simulation results, high serotonergic fiber densities can decrease more gradually away from the border; however, this property can be mirrored in simulated densities by changing the convexity of the transforming function (effectively, by deciding at which density value intensities saturate and become virtually indistinguishable).

The hippocampal complex was modeled as a featureless, unfolded medial pallial region (Butler and Hodos, 2005; Striedter and Northcutt, 2020). This representation was deliberately inaccurate, to avoid the 3D-reconstruction of a complex, layered structure (with the correctly placed entrance and exit zones for fibers). Unsurprisingly, the low simulated density in this area (Fig. 3C) deviated strongly from the generally high density of serotonergic fibers in the hippocampus (Awasthi et al., 2020). However, if the folding had been simulated, a high fiber density would have been observed in the superficial layer of this *cortical* structure (the archicortex). In the hippocampus proper, this superficial layer corresponds to the stratum lacunosum moleculare, which has the highest serotonergic fiber density among all layers (Vertes et al., 1999; Nazzi et al., 2019; Awasthi et al., 2020). Likewise, the molecular layer of the mouse dentate gyrus has been reported to have the highest serotonergic fiber density among all layers (Awasthi et al., 2020). A slightly different pattern has been reported in another study that has focused only on the fibers originating in the median raphe nucleus of the rat (Vertes et al., 1999). This discrepancy may be due to an incomplete labeling of fibers with the tracer.

The mediolateral fiber-density profile in the septum (Fig. 3B) was remarkably consistent with the reported serotonergic densities in this region (Awasthi et al., 2020), with a high peak in the medial septum, a deep trough in the medial part of the lateral septum, and another peak in the lateral part of the lateral septum, at the edge of the lateral ventricle (see Fig. 11 of Vertes et al. (1999)). This close match may be coincidental, but it may also reflect the potential of the proposed approach. The observed profile cannot be deduced from a single coronal section and it may also have a developmental component, since in the adult brain this region is dorsally limited by the thick band of the corpus callosum (which was not modeled in the used 3D-shape).

The current model cannot capture local variations in serotonergic fiber densities that are not accounted for by the geometry of the borders. For example, the cortical layers below layer I often do not follow a descending gradient and can produce local density peaks deeper in the tissue, such in layers IV or V (Linley et al., 2013; Awasthi et al., 2020). Likewise, individual nuclei in the amygdala complex can be located close to one another but differ strongly in their serotonergic fiber densities (Awasthi et al., 2020). This local variability may be due to local biological factors that control the growth and branching of fibers. These discrepancies do not signal fundamental limitations of the proposed model which assumed that the interior of the 3Dshape was a uniform medium, with no spatial heterogeneities. In reality, such heterogeneities are always present in natural neural tissue and can be easily observed even in unstained preparations (Fig. 1). They include regional variations in neuron packing densities (Erö et al., 2018; Keller et al., 2018), extracellular space (Hrabetova et al., 2018), viscoelasticity (Cherstvy et al., 2019; Antonovaitė et al., 2021), and other variables. It is currently unclear which of these factors can affect the resultant densities of simulated fibers; for example, our limited analysis of rFBM-fibers in 2D-shapes with densely packed cells (small obstacles) produced results similar to those in corresponding shapes with no cells (Janušonis et al., 2020). The incorporation of spatial heterogeneities requires advances in the theory of FBM, where a non-constant *H* poses challenges in mathematical specifications of the process. We are currently developing new theoretical models that can overcome these difficulties (Wang et al., 2023). It is worth noting that the densities of glial cells show surprisingly little regional variability, and even little variability across mammalian species (Herculano-Houzel, 2014; Dos Santos et al., 2020).

The primary aim of the simulation was not to fully replicate the regional densities of serotonergic fibers but rather to examine to what extent these densities can be explained by the intrinsic stochastic properties of the fibers, as they interact with the basic geometry of the brain. We note that different axon types may lie at different points along the stochastic-deterministic continuum. For example, the trajectories of the retinogeniculate and corticospinal axons can be considered *strongly deterministic*. In contrast, serotonergic fibers appear to be *strongly stochastic*; in the adult brain they typically produce highly tortuous trajectories (sometimes with complete loops), can meet at any angles (Janušonis et al., 2019), and do not fasciculate. The stochasticity of axon growth has been acknowledged by a small number of other studies (Katz et al., 1984; Maskery and Shinbrot, 2005; Betz et al., 2006; Yurchenko et al., 2019), but thus far these approaches have not led to major applications in neuroscience. The fundamental problem of the self-organization of serotonergic fiber densities may motivate this line of research, which can be enriched with other theoretical insights, such Brownian ratchet theory and Wiener sausage-like constructs.

At the qualitative level, the rFBM-model of serotonergic fibers can make intriguing predictions. For example, the accumulation of fibers at a border is likely to be higher if the border has a high local curvature. We have demonstrated this phenomenon in 2D-shapes (Janušonis et al., 2020; Vojta et al., 2020). Since the size of neurons and axons generally does not scale with the brain, it implies that larger brains might have relatively lower serotonergic fiber densities at the surface – unless the curvature is restored with gyrification. It also implies that the shape of the brain, whether molded by evolution (Striedter and Northcutt, 2020), artificial selection, or artificial cranial deformation (Meiklejohn et al., 1992) might affect the distribution of serotonergic fibers, with potential implications for regional neuroplasticity.

Irrespective of the exact specification of the used stochastic process, this study highlights the possibility that serotonergic fiber densities are never truly local. In order to reach a specific target, fibers have to traverse other regions, introducing spatial correlations. Also, the absence of physical tissue or the emergence of obstacles in adjacent planes prevents fibers from advancing in this direction and may give the impression of an external guiding force when viewed in only one plane.

The strong stochasticity of serotonergic fibers does not exclude other factors that may guide their interaction with the environment and with one another. These factors include brain-derived neurotrophic factor (BDNF) (Mamounas et al., 1995), the growth factor S100β (Sodhi and Sanders-Bush, 2004), GAP-43 (Donovan et al., 2002), the microtubule-associated STOP proteins (Fournet et al., 2010), and protocadherins (Katori et al., 2009; Chen et al., 2017; Katori et al., 2017). In addition, 5-HT itself may affect the growth of serotonergic fibers (Whitaker-Azmitia, 2001; Migliarini et al., 2013), but these effects are neuroanatomically subtle (Lesch et al., 2012; Montalbano et al., 2015; Mosienko et al., 2015). They can also be masked by the availability of placenta-derived 5-HT in fetal brain development (Bonnin and Levitt, 2011). Interestingly, serotonergic fibers themselves may guide postnatal neuroblast migration (García-González et al., 2017).

Alterations in regional serotonergic fiber densities have been associated with a number of mental disorders and conditions, such as autism spectrum disorder (Azmitia et al., 2011), epilepsy (Maia et al., 2019), major depressive disorder (Numasawa et al., 2017), social isolation (Keesom et al., 2018), and the abuse of 3,4-methylenedioxy-methamphetamine (MDMA, Ecstasy) (Adori et al., 2011). In the healthy brain, 5-HT signaling plays major roles in the sleepwake cycle (Brown et al., 2012) and reward circuits (Liu et al., 2020). It suggests that predictive models of the self-organization and dynamics of serotonergic fibers may advance not only basic neuroscience but also find applications in the biomedical field.

## 6. Author Contributions

SJ proposed that the self-organization of serotonergic densities can be modeled with FBM-paths, produced the embryonic brain section series (with JHH), wrote the Mathematica scripts to convert raw section images into the simulation-compatible format, and wrote the first draft of the manuscript. TV performed all supercomputing simulations. RM contributed to the theoretical development of the used approaches. SJ, RM, and TV are the Principal Investigators of their respective research programs.

## 8. Acknowledgements

We acknowledge Dr. Kasie Mays’ (UCSB) advice on published serotonergic density maps and Parsa Madinei’s (UCSB) assistance with histology.

